# The D84G mutation in STIM1 causes nuclear envelope dysfunction and myopathy in mice

**DOI:** 10.1101/2023.05.03.539279

**Authors:** Victoria Bryson, Chaojian Wang, Zirui Zhou, Kavisha Singh, Noah Volin, Eda Yildirim, Paul Rosenberg

**Author notes:** these authors contributed equally to this work (Victoria Bryson, Chaojian Wang).

## Abstract

Stromal interaction molecule 1 (STIM1) is a Ca^2+^ sensor located in the sarcoplasmic reticulum (SR) of skeletal muscle where it is best known for its role in store operated Ca^2+^ entry (SOCE). Genetic syndromes resulting from STIM1 mutations are recognized as a cause of muscle weakness and atrophy. Here, we focus on a gain of function mutation that occurs in humans and mice (STIM1^+/D84G^ mice) where muscles exhibit constitutive SOCE. Unexpectedly, this constitutive SOCE did not affect global Ca^2+^ transients, SR Ca^2+^ content or excitation contraction coupling (ECC) and was therefore unlikely to underlie the reduced muscle mass and weakness observed in these mice. Instead, we demonstrate that the presence of D84G STIM1 in the nuclear envelope of STIM1^+/D84G^ muscle disrupts nuclear-cytosolic coupling causing severe derangement in nuclear architecture, DNA damage, and altered lamina A associated gene expression. Functionally, we found D84G STIM1 reduced the transfer of Ca^2+^ from the cytosol to the nucleus in myoblasts resulting in a reduction of [Ca^2+^]_N_. Taken together, we propose a novel role for STIM1 in the nuclear envelope that links Ca^2+^ signaling to nuclear stability in skeletal muscle.

## Introduction

Stromal interaction molecule 1 (STIM1) is a transmembrane protein located in the sarcoplasmic reticulum (SR) of skeletal muscle where it senses Ca^2+^ stores and activates Orai channels as part of store operated Ca^2+^ entry (SOCE). SOCE in muscle fibers is required for Ca^2+^ homeostasis and to sustain Ca^2+^ transients during neuromuscular activity(1). Loss of function mutations in the human and mouse genes for STIM1 and Orai1 cause reduced muscle growth and weakness, in addition to severe combined immune deficiency(2-4). Gain of function mutations in STIM1 and Orai1 are increasingly recognized as a genetic syndrome that includes hyposplenism, platelet bleeding diathesis, and tubular aggregate myopathy (TAM1)(5). Tubular aggregates (TA) form in specific regions of the SR membrane that contain STIM1, Calsequestrin and SERCA1 but exclude the ryanodine receptor (RYR1). While TAs are associated with STIM1 mutations, it remains to be determined if they are required for the STIM1 myopathy. In fact, various mouse models of the STIM1 mutations fail to manifest TAs but do experience weakness(6-10).

In skeletal muscle, the SR is comprised of highly specialized regions that include the junctional cisternae (jSR), longitudinal SR (lSR) and the perinuclear membrane (PNS). The ISR houses the RYR1, is located adjacent the A-I bands of the sarcomere, and is the site of Ca^2+^ release during excitation contraction coupling (ECC). The longitudinal SR encompasses the fenestrated membrane near the Z-line that is enriched in SERCA1 and functions in Ca^2+^ store refilling, protein synthesis and protein secretion. STIM1 is organized into operational domains of the SR membrane that include the longitudinal SR and terminal cisternae. STIM1 in the terminal cisternae interacts with Orai1 in the T-tubular membrane to activate SOCE. We recently showed that STIM1 is tethered in the longitudinal SR by the type III intermediate filament (IF) desmin to regulate SERCA1 and Ca^2+^ signaling in the muscle fibers(11). Desmin anchors STIM1 to the Z-disc of muscle fibers and regulates SOCE and prevents Ca^2+^ overload by facilitating store refilling by SERCA1 pumps.

In contrast to our understanding of STIM1 in the SR membrane, far less is known about the role STIM1 has in the nuclear membrane (12). STIM1 can be detected in the outer nuclear membrane and as part of the nuclear reticulum. We hypothesized that STIM1 has an essential role in nuclear-cytosolic connectivity and interacts with proteins of the linker of nucleoskeleton and cytoskeleton (LINC) complex. The LINC complex is composed of several proteins that span the nuclear envelope and include Lamin A/C in the nuclear lamina, Sad1 and UNC84 domain containing proteins (SUN1/2) of the inner nuclear membrane (INM) and Kash/Nesprin proteins of the outer nuclear membrane (ONM)(13). Compromise of the nuclear-cytosolic connectivity can influence nuclear morphology, induce apoptosis, and influence gene expression, as occurs in Emery Dreifuss Muscular Dystrophy (EDMD) and other laminopathies(14).

In the present work, we used a mouse model bearing a mutation in the EF hand (D84G) of STIM1 to explore the role of STIM1 in the nuclear membrane *in vivo* (15, 16). We show that STIM1^+/D84G^ mice exhibit constitutively activated SOCE but muscle fibers are spared the consequence of Ca^2+^ overload due to remodeling of Ca^2+^ handling apparatus in these muscles. Intriguingly, we found that changes in the nuclear morphology that resemble changes observed in myopathies associated with laminopathies. These findings offer novel insight to the pathogenesis of STIM1-myopathies and demonstrate a previously unrecognized role for STIM1 in the nuclear envelope, gene expression and DNA damage.

## Results

### Reduced muscle mass in STIM1^+/D84G^ mice

STIM1^+/D84G^ mice are fertile and exhibit no excess mortality compared to WT littermates. We did however observe a significant reduction in body weight in a cohort of STIM1^+/D84G^ mice (Figure 1A). At 6-months of age, a reduction in muscle mass was observed for specific muscles including tibialis anterior (TA) and extensor digitorum longis (EDL) muscles (Figure 1B). Mass of soleus muscles (Sol) were not different (Figure 1B). Tibial length did not differ for STIM1^+/D84G^ and WT mice (data not shown). These data suggest that the loss of muscle mass for STIM1^+/D84G^ mice selectively involves muscles enriched in fast glycolytic fibers whereas muscles enriched in slow oxidative fibers maintain muscle mass.

**Figure 1:**
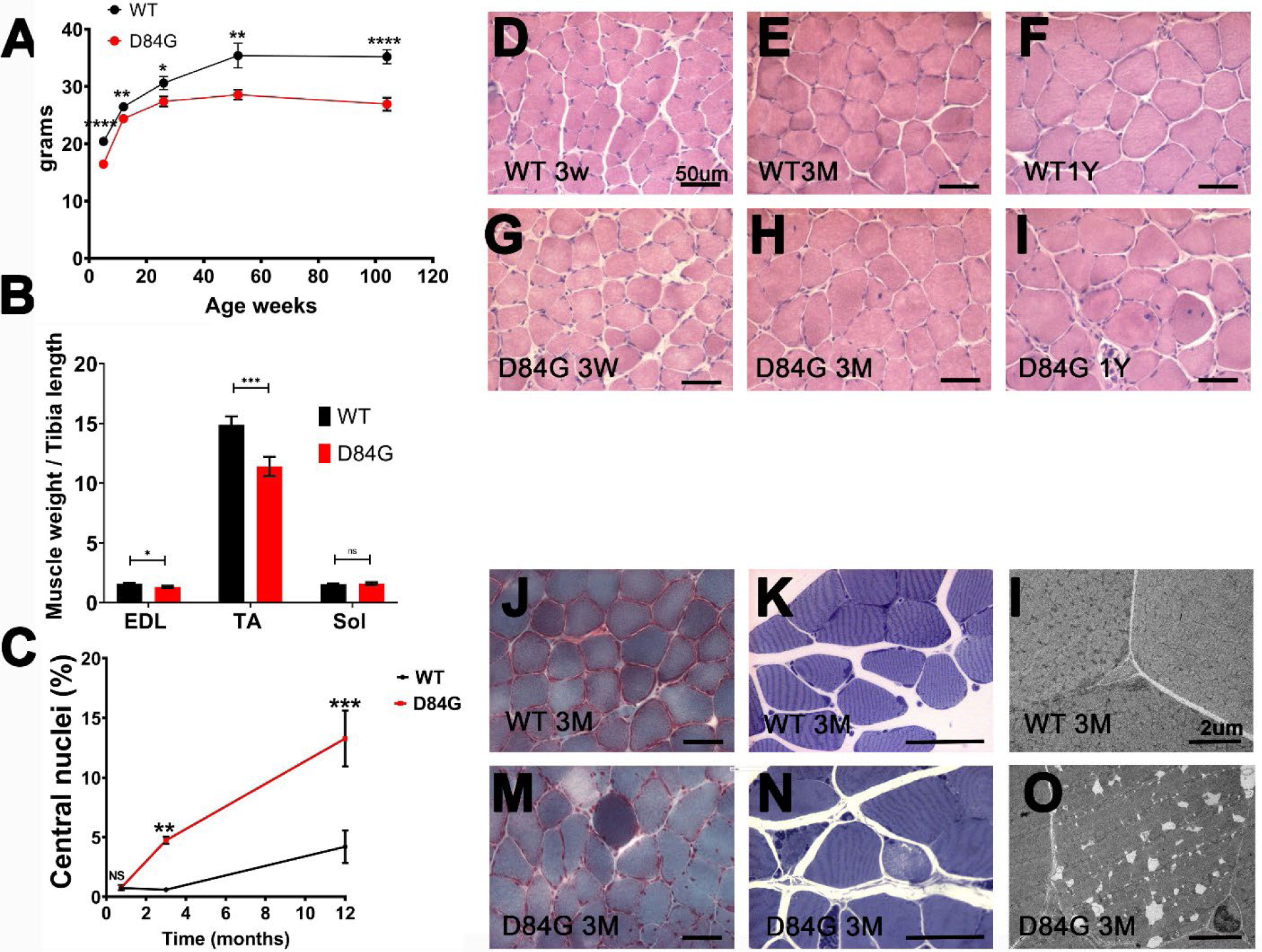
Characterization of STIM1^+/D84G^ muscle. (A) Mouse weight with time (n=6-9 mice per genotype) (B) Muscle weight to tibia length ratio of extensor digitorum longis (EDL), soleus (Sol) and tibialis anterior (TA) muscle of 6 months old WT and D84G mice (n=6 per genotype) (C) Quantification of central nuclei percentage in the soleus muscle from WT and D84G mice (n=3). (D-I) Hematoxylin and Eosin staining of 14um cryosections of soleus muscle from WT and D84G mice (D) WT 3 weeks (G) D84G 3 weeks (E) WT 3 months (H) D84G 3 months (F) WT 1 year (I) D84G 1 year (n=3). (J and M) Gamori trichrome staining of 3-month soleus muscle sections (n>3) (J) WT (M) D84G. (K and N) Toluidine blue staining of resin thin sections of TA muscle (K) WT and (N) D84G (L and O) Transmission electron microscopy (TEM) micrographs of ultrathin TA (tibialis anterior) muscle sections of (L) WT and D84G (O) (n>3). Values are mean ± standard error. Two-tailed Student’s *t*-test, n ≥ 4-6 independent experiments. NS, not significant (*p* > 0.05). *p<0.05, **p<0.01, ***p<0.001, ****p<0.0001

### Histopathology of the STIM1^+/D84G^ muscle

Histopathology for STIM1^+/D84G^ skeletal muscles, evaluated by Hematoxylin and Eosin stained sections, revealed a progressive increase in the number of central nuclei over one year (Figure 1C-I). GÖmÖri staining of muscle sections from STIM1^+/D84G^ mice did not reveal changes to the SR that are typically seen for tubular aggregates, however muscle fibers were variable in size and color ranging from pale necrotic fibers to intensely stained pink fibers, findings consistent with excess contractile damage (Figure 1J and M). Toluidine blue stained sections of 3-month old STIM1^+/D84G^ mice identified abnormal shaped and pale fibers, necrotic fibers and expansion of the extracellular matrix; non-specific findings often seen in skeletal myopathies (Figure 1K and N). Transmission electron microscopy (TEM) micrographs of TA muscles from 3-month old WT and STIM1^+/D84G^ mice were examined. Structural changes noted from muscle sections of STIM1^+/D84G^ mice included marked dilation of the terminal cisternae of the SR and distortion of the longitudinal SR that resembles SR stacking in tibialis anterior and soleus muscles (Figure 1L and O). Tubular aggregates (TA) were rarely detected in the muscles of STIM1^+/D84G^ mice. Swollen mitochondria were also detected in areas of myofibril contractures (data not shown). These findings resemble the steps described for TA development in aged muscles where SR dilation precedes TA formation (17).

### Muscle-specific gene expression profiling of D84G mice

To understand the differences in gene expression between muscles from WT and STIM1^+/D84G^ mice, we performed RNA sequencing for gastrocnemius muscles isolated from six-month old mice. The volcano plots display differentially expressed genes (DEG) from STIM1^+/D84G^ mice that include 15,434 transcript reads. Of the total transcripts, 1,543 DEGs (∼10%) were upregulated and only 116 DEGs (∼0.8%) were downregulated based on statistical cutoff of normalized enrichment score (NES>2.5) and FDR<0.05 (Figure 2A-B, Supplemental Data Table 1).

**Figure 2:**
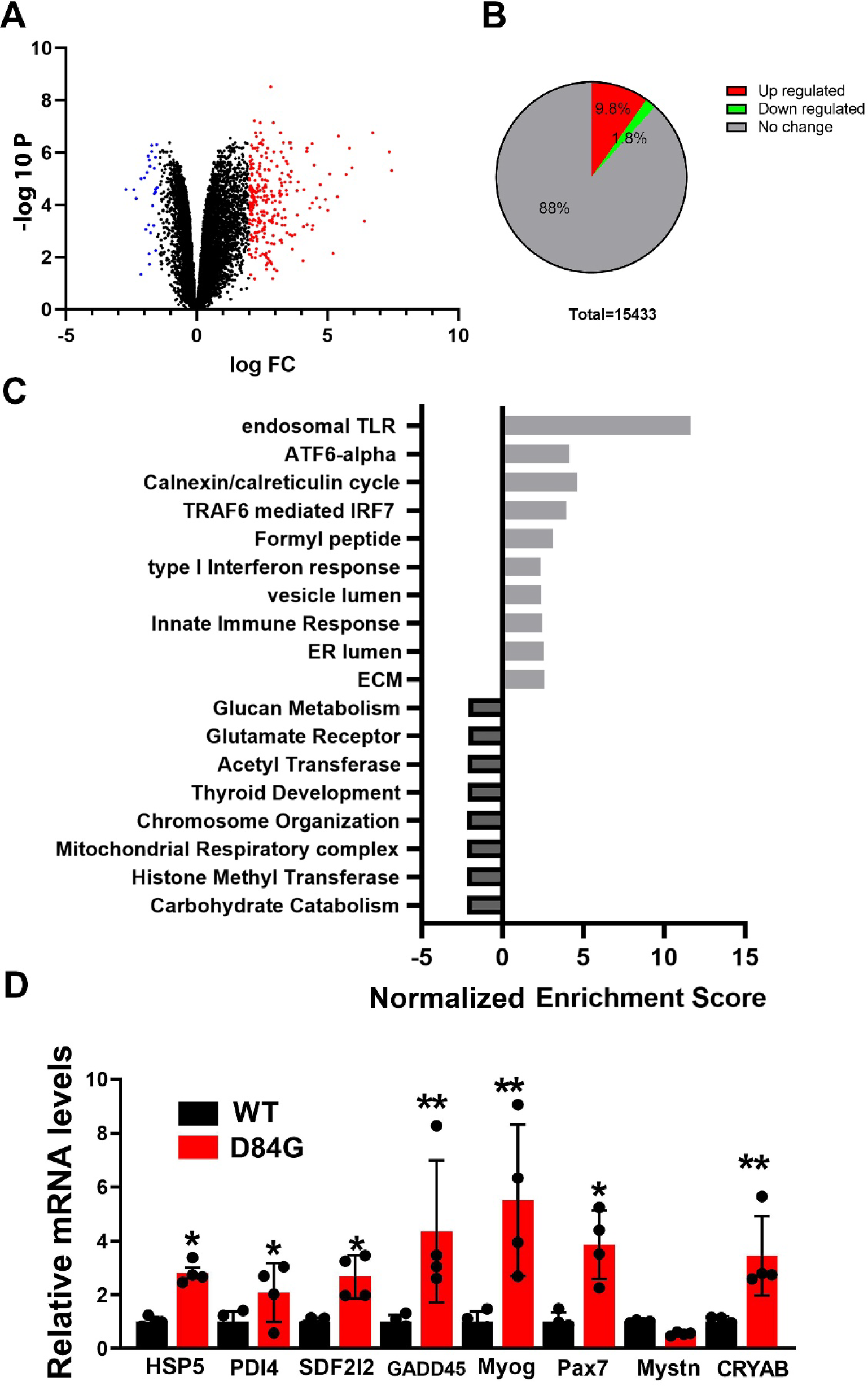
RNA sequencing for gastrocnemius muscles from 6-month old WT and STIM1^+/D84G^ mice. mRNA from 6-month old WT (n=4) and STIM1^+/D84G^ (n=4) was prepared from gastrocnemius muscles. (A) Volcano plots for differentially expressed genes (DEGs) from WT and STIM1^+/D84G^ mice. Red dots represent upregulated and blue dots represent downregulated genes. (B) Graphic representation for DEGs upregulated and down regulated. (C) Pathway analysis for DEGs was performed using GO pathway analysis. The top pathways for upregulated DEGs (black) and downregulated DEGs (gray) are shown. (D) Quantification of ER stress genes expression from STIM1^+/D84G^ muscles using rt-PCR. Values are mean ± standard deviation. Relative mRNA levels were normalized using GAPDH. Two-tailed Student’s *t*-test, *n* ≥ 4 independent experiments. NS, not significant (*p* > 0.05). *p<0.05, **p<0.01

### Disease pathways in skeletal muscle of STIM1^+/D84G^ mice

Gene Ontology (GO) Analysis identified significant changes in several biologic processes and pathways for the upregulated DEGs that associate with inflammation (TLR, IFN signaling, type I immune response), ER stress (ATF-6, calnexin, ER lumen), myogenesis and extracellular matrix (Figure 2C). Involvement of the ER stress and unfolded protein pathways was validated by quantifying the mRNA by qPCR for Heat Shock Protein Family A (HSP70) Member 5 (HSPA5), Protein Disulfide Isomerase Family A Member 4 (PDIA4), Growth Arrest And DNA Damage Inducible Alpha (GADD45) and Stromal Cell Derived Factor 2 Like 1 (SDF2L1). ER-stress pathways sense accumulation of the misfolded protein and activate transcriptional pathways that upregulate chaperones in the ER to promote refolding (Figure 2D). Myogenic factors were also upregulated include Myogenin (MYOG), Paired Box 7 (PAX7), Myostatin (MSTN) and αβ-crystallin (CRYAB).

Based on the DEGs downregulated in the muscles of STIM1^+/D84G^ mice, GO enrichment tools identified signaling pathways associated with carbohydrate catabolism, histone methyltransferase activity, mitochondrial oxidative metabolism, DNA repair, chromosome organization and chromatin modification (Figure 2C). These pathways are frequently linked to chromatin organization, muscle differentiation and mitochondrial biogenesis. Taken together, marked changes in the transcriptome of muscles from STIM1^+/D84G^ mice lead to changes in pathways related to ER stress and immune response are upregulated; whereas pathways associated with chromatin regulation were down regulated.

### Spontaneous Ca^2+^ entry into the D84G muscle fibers

The N-terminus of STIM1 extends into the SR lumen where EF hands senses depletion of luminal Ca^2+^ stores(18). The D84G mutation in STIM1 disrupts the globular compact structure created by the EF hand and SAM domains and thereby reducing Ca^2+^ bound to it. As shown by others, this mutant STIM1 associates with Orai1 channels in the absence of store depletion and confers spontaneous Ca^2+^ entry(19). To examine the effect of D84G STIM1 on Ca^2+^ entry in FDB muscle fibers, we performed Manganese (Mn^2+^) quenching assays by exciting Fura-2AM (360 nm) at the isosbestic wavelength. Manganese (1.8 mM) application to the cellular bath solution is detected as loss of the emission of Fura-2 signal and thereby represents cationic flux across the sarcolemma. This enables the separation of Ca^2+^ entry and Ca^2+^ release and is known to be linearly related to SOCE. Addition of Mn^2+^ (1.8 mM) to D84G fibers reduced the fluorescence signal in the absence of store depletion that was greater than WT fibers, consistent with increased spontaneous Ca^2+^ entry in D84G muscle (Figure 3A-B). SOCE is activated during neuromuscular activity of WT fibers by both tonic and phasic patterns (20). Electrical field stimulation (EFS) of WT fibers led to greater rate of Fura-2 quenching compared to the rate in the absence of EFS within the same fiber; findings that are consistent with SOCE phasic activation (Figure 3A-B). Importantly, no differences in the rate of Fura-2 quenching was detected for the EFS-stimulated WT and STIM1^+/D84G^ fibers (Figure 3A and C). These data suggest that D84G STIM1 can spontaneously activate Orai1 channels in the T-tubular membrane. Because the STIM1^+/D84G^ fibers also contain the WT STIM1, STIM1 Ca^2+^ sensor function was intact as SOCE was activated by EFS. Despite the spontaneous Ca^2+^ entry in the D84G fibers, no differences in basal Ca^2+^ levels (Figure 3D). In addition, Ca^2+^ release from internal stores induced by caffeine (RYR1 activator) and SERCA inhibition by cyclopiazonic acid (CPA) were detected for STIM1^D84G^ and WT mice (Figure 3E-F).

**Figure 3:**
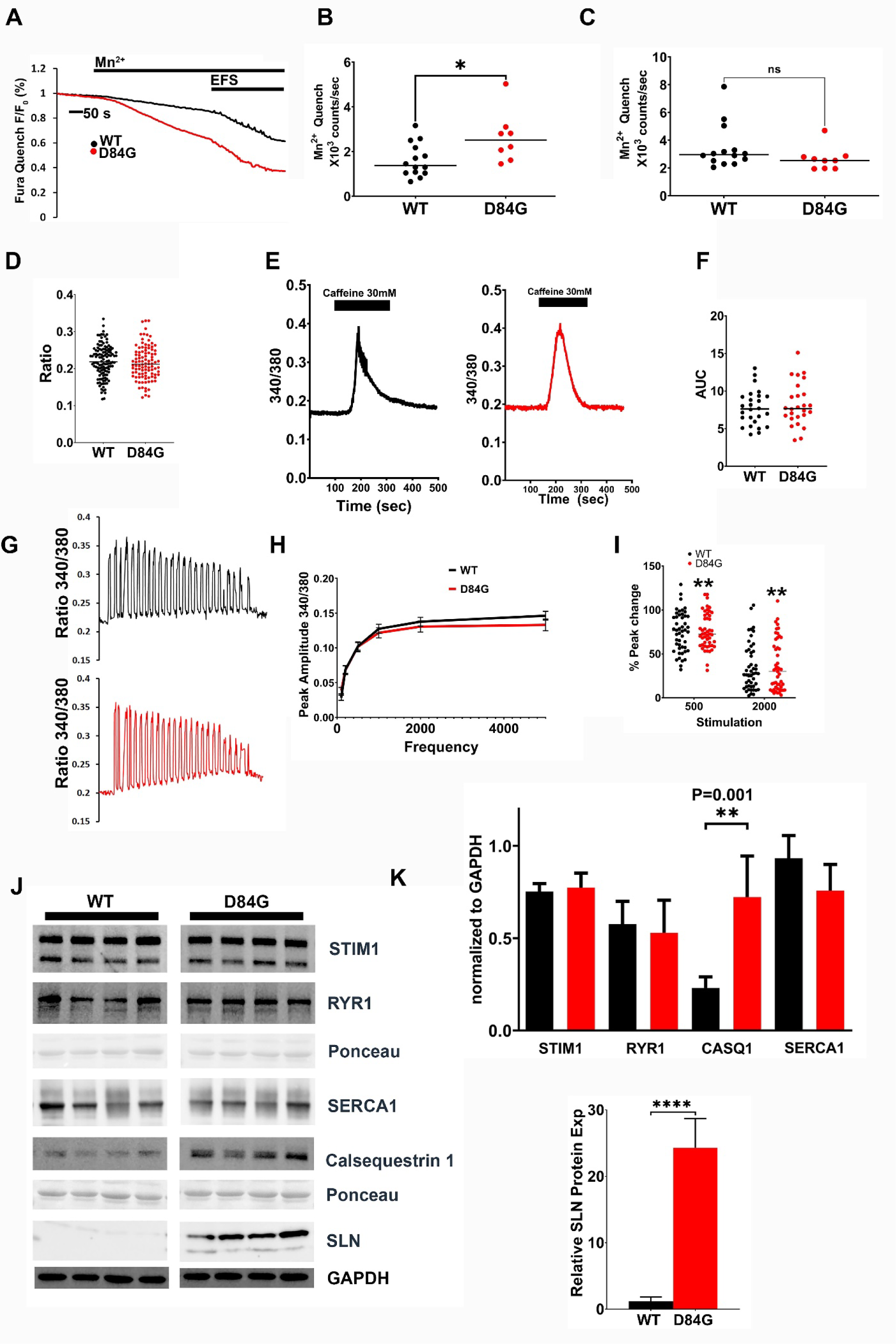
Ca^2+^ signaling in the D84G mutant mice. (A-C) Manganese (Mn^2+^) quench assays were performed on Fura-2 loaded FDB fibers to quantitate SOCE from WT and STIM1^+/D84G^ mice (n=3 mice, 30 fibers)(A). Mn^2+^ quenching rate was considered spontaneous after 150 s after Mn^2+^ addition (B). The Mn^2+^ quenching following electrical field stimulation (EFS) was determined but was not different (C). (D) Basal Ca^2+^ levels measured by the Fura-2 method were not different between WT and STIM1^+/D84G^ fibers. (E-F) Ca^2+^ release evoked by caffeine from Fura-4F loaded FDB fibers (E). Area under the Curve (AUC) (F) did not differ between WT and STIM1^+/D84G^ fibers (n=3 mice, 30 fibers per genotype). (G-I) EFS Ca^2+^ transients from Fura-4F loaded FDB muscle fibers were similar (n=4 mice with 50 fibers per genotype) (G). Graphical representation of peak amplitude and frequency (H) demonstrate no significant change to the peak Ca^2+^ release per EFS but a significant elevation in interstimulus Ca^2+^ at 500 ms and 2 ms from STIM1^+/D84G^ fibers n=50 fibers per genotype (3 mice) (I). **p<0.01. (J-K) Western blot analyses of Ca^2+^ handling proteins from 6-month WT and STIM1^+/D84G^ mice (n=5 per genotype). Lysates from Gastrocnemius muscles were prepared from WT and STIM1^+/D84G^ mice. Antibodies for STIM1, RYR1, SERCA1, Calsequestrin (CASQ) (left) and sarcolipin (SLN) (right) were used to quantify protein expression vs GAPDH. Values are mean ± standard deviation. Relative protein levels were normalized using GAPDH. Two-tailed Student’s *t*-test, *n* ≥ 4 independent experiments. NS, not significant (*p* > 0.05) **p<0.001. ****p<0.0001.

To understand how the D84G STIM1 mutation influences EFS-evoked Ca^2+^ transients, we measured transients from Fura-4-loaded FDB fibers (21). No difference in peak amplitude of the Ca^2+^ transients was detected over a range of stimulation frequencies (1-50 Hz) for WT and STIM1^+/D84G^ fibers (Figure 3G-H). However, analysis of the inter-stimulus nadir of Ca^2+^ revealed an increase in the Ca^2+^ for the STIM1^+/D84G^ fibers (Figure 3G and I). Specifically, the change in Ca^2+^ was apparent after 500 s and 2000 s of EFS. These data may represent the spontaneous SOCE that persists in the stimulated fiber. From these data it appears the presence of D84G STIM1 in the SR did not significantly alter the Ca^2+^ transients. We therefore scrutinized RNA-seq data from STIM1^+/D84G^ muscle (Figure 2) for changes in expression for factors that regulate Ca^2+^ handling proteins. Sarcolipin (SLN), an endogenous muscle specific inhibitor of the SERCA1 pump, was among the most dramatically upregulated mRNA (∼40-fold) in STIM1^+/D84G^ muscle and the corresponding increase in SLN protein levels (Figure 3J-K)(22). STIM1, SERCA1 and RYR1 expression were unchanged for the gastrocnemius muscles of STIM1^+/D84G^ mice (Figure 3J-K). In contrast, the SR-Ca^2+^ buffering protein calsequestrin 1 (CASQ1) was down regulated in the RNA-seq data but the level of CASQ1 protein was significantly increased in STIM1^+/D84G^ muscle lysates. Taken together these changes in Ca^2+^ handling proteins, such as SLN and CASQ1, likely represent an important adaptive mechanism available to muscle fibers that protect fibers against Ca^2+^ store overload that would otherwise be expected when SOCE is constitutive.

### Impaired exercise capacity for D84G mice

To assess general locomotor activity for STIM1 mutant mice, we measured open field exploratory behavior for WT and STIM1^+/D84G^ mice. STIM1^+/D84G^ mice exhibited a reduction in the distance explored that was significantly less compared to WT counterparts (Figure 1A). WT mice progressively explore their environment over the entire 15-minute interval. However, there was little cumulative movement for the STIM1^+/D84G^ mice. These data identify a progressive decline locomotor activity for STIM1^+/D84G^ mice over time compared to WT mice. We next assessed maximal exercise capacity for 3-month old STIM1^+/D84G^ mice *in vivo* using a standard treadmill running protocol (Figure 4B). The significant reduction in time to exhaustion and distance run for the STIM1^+/D84G^ mice demonstrate that these mice have limited exercise capacity compared to WT mice (Figure 4A). Grip strength was also significantly reduced for 6-month old STIM1^+/D84G^ mice compared to WT mice (Figure 4C). Despite relatively normal Ca^2+^ transients, the STIM1^+/D84G^ mice experience progressive decline in locomotor function and reduced exercise capacity.

**Figure 4:**
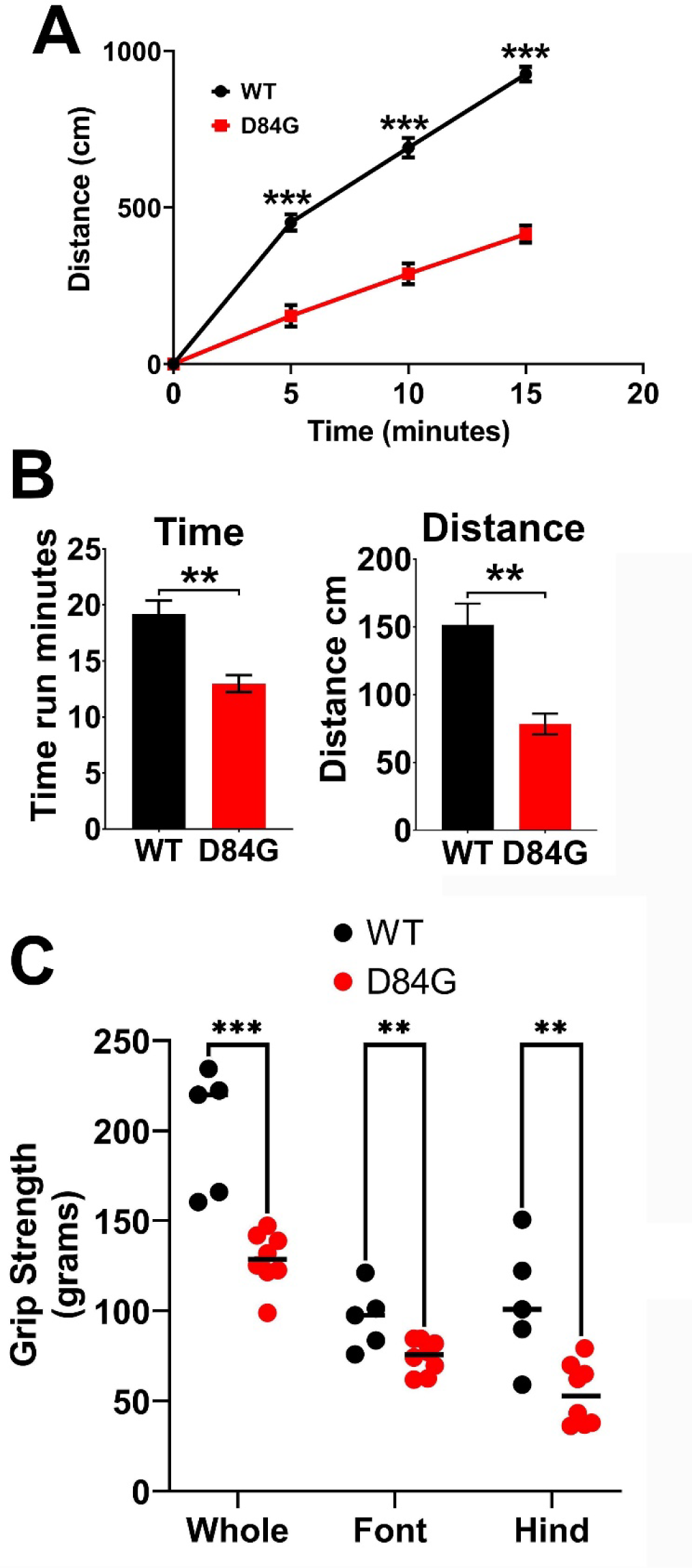
Muscle performance is reduced in the STIM1^+/D84G^ mice. (A) Spontaneous locomotion was quantified using an exploratory open field platform. Recordings of movement every 5-minutes were quantified for WT and STIM1^+/D84G^ mice. Mice were measured at 1-year (n=5 mice) for each genotype. (B) Treadmill running distance (left) and times (right) for 3-month old WT (n=6) and STIM1^+/D84G^ mice (n=6). (C) Grip strength was measured for front, rear and whole body (n=6). Values are mean ± standard deviation. Two-tailed Student’s *t*-test, n ≥ 6 independent experiments. NS, not significant (*p* > 0.05). *p<0.05. **p<0.01, ***P<0.001

### STIM1 in the nuclear membrane

Given the changes to muscle mass and altered SR ultrastructure in muscles of STIM1^+/D84G^ mice, we next considered that the mutant D84G STIM1 was not properly targeted in the SR. FDB fibers from WT mice immunostained with STIM1 antibodies and DAPI to label nuclei demonstrate STIM1 in the terminal cisternae, the longitudinal SR near the Z-line and throughout the nuclear membrane, as previously shown(11). STIM1 is enriched in the nuclear reticulum (NR) which is a specialized NE invaginations (NEI) that extend from the cytosol into the nucleoplasm (Figure 5A). Confirmation of STIM1 in the nuclear membrane was obtained by transmission electron microscopy (TEM) of muscles taken from STIM1 reporter mice (STIM1^+/LacZ^ mice) where one copy of STIM1 is fused to beta-galactosidase(4). The STIM1-LacZ appears as a black precipitate and is clearly expressed in the nuclear envelope (Figure 5B).

**Figure 5:**
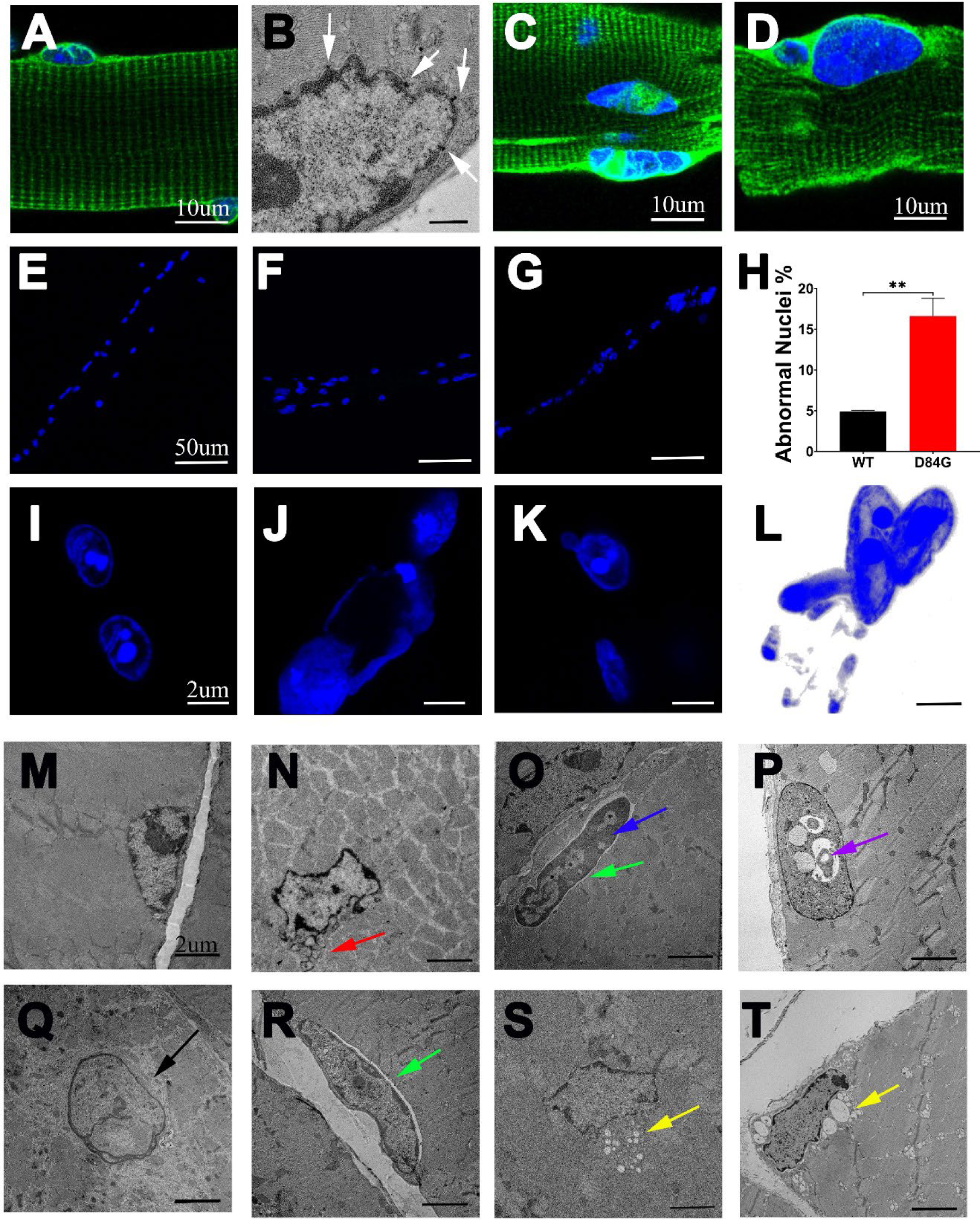
Nuclear abnormalities in muscle of STIM1^+/D84G^ mice. (A) Immunostaining for STIM1 (green) and nuclei (DAPI, blue) in WT mice, scale 10um (n>6). (B) Electron microscopy showing expression of STIM LacZ (white arrows) in the nuclear envelope of STIM1^gt/+^ mice as detected by X-gal staining, scale 500nm (n>4). (C-D) STIM1 expression in D84G mice, scale 10um. (E-L) Nuclear abnormalities of D84G mice as demonstrated by DAPI staining (blue) (n>6). (E-G) Nuclei along the length of the fiber in WT (E) and D84G mice (F-G), scale 50um. (H) Quantification of abnormal nuclei (large, clumped, lobular or fragmented) (%) in WT and D84G mice. Values are mean ± standard deviation. Two-tailed Student’s *t*-test, *n* ≥ 6 independent experiments. *p<0.05. (I-L) High power confocal images of WT (I) and D84G (J-K) DAPI stained nuclei, scale 2um (L) is a Maximum image projection through a D84G nucleus, scale 2um. (M-R) Transmission electron micrographs of nuclei in the TA muscle of WT (M) and D84G mice (N-R) (n>3). (N) D84G nucleus with fragmented micronuclei, red arrows (O and R); D84G nuclei with dilated perinuclear space, green arrow (O); D84G with condensed chromatin, blue arrow (P); Vacuolated nucleus in D84G mice, purple arrow (Q); pale fading nucleus suggesting karyolysis and nuclear rupture (S and T).

In STIM1^+/D84G^ muscle fibers, STIM1 was detected in the longitudinal SR, in a striated pattern typical for the SR (Figure 5C and D). Nuclear staining in STIM1^+/D84G^ muscle fibers resembled that seen in the WT fibers for many nuclei. However, a number of nuclei were observed with abnormal STIM1 staining, these altered patterns were associated with altered nuclear morphology. STIM1 staining in the NE of STIM1^+/D84G^ muscle fibers often appeared distorted and occupied a much broader region of the NE. Here, STIM1 was detected throughout the NE and can be seen bisecting the nucleus unevenly from what appears to be abnormal NEI as DAPI (a marker for DNA) was absent from this region (Figure 5C and D). Nuclear morphology varied enormously in STIM1^+/D84G^ fibers ranging from small micronuclei to massive nuclei with multiple lobes (Figure 5 E-G, J-L). WT nuclei are always positioned underneath the sarcolemma, whereas D84G nuclei can be seen in the middle of fiber often displacing myofibrils. We quantified nuclear abnormalities in both genotypes and found ∼16% of nuclei from STIM1^+/D84G^ fibers appeared clumped, aggregated, misshapen or broken down into micronuclei. In contrast only ∼5% of WT nuclei displayed these characteristics (Figures 5E-L). These data suggest that the presence of the mutant STIM1 in these fibers compromised the structure and integrity of the NE. Given the changes to nuclear shape of STIM1^+/D84G^ fibers, we used TEM to better assess the nuclear architecture of skeletal muscle (Figure 5M-T). We found evidence for nuclear rupture and herniation, dense chromatin accumulation and markedly dilated nuclear invaginations. Importantly, the perinuclear space was also found to be dilated for myonuclei from STIM1^+/D84G^ muscles. Finally, we were able demonstrate connections, albeit rare, between the TAs in the SR and the dilated PNS in some fibers (Figure 5O). These data show that constitutively active STIM1 mutations disrupt nuclear structures which contribute to pathogenesis for the STIM1 myopathy.

### D84G STIM1 destabilizes the LINC complex and nuclear lamina

We next considered recently published database by Gu et al. where proximity BioID proteomics was performed using STIM1 as one of many baits in HEK293 cells (23). Of the 173 proteins identified to be in close proximity of STIM1 with this assay are proteins located at the ER, golgi and NE. We used NIH DAVID Bioinformatics Resources (https://david.ncifcrf.gov/home.jsp) to identify signaling pathways and disease terms associated with this group of proteins and found terms including nuclear cytosolic transport and Emery Dreifuss Muscular Dystrophy (EDMD) were significantly linked to STIM1 proximity (Supplemental Data Table 1)(23, 24). We therefore hypothesized that components of the nuclear lamina, which is the meshwork of intermediate filaments that form part of the NE, would be altered in the muscle fibers of STIM1^+/D84G^ mice (25). LMNB1 and SUN2 expression were unchanged in nuclear extracts of STIM1^+/D84G^ muscle (Figure 6A, Supplemental Figure 2A). LMNA/C can be detected as two bands on immunoblots for WT muscle consistent with the LMNA (74kD) and LMNC (63kD) isoforms. A significant increase in the level of LMNA isoform was apparent in the nuclear extracts of STIM1^+/D84G^ mice whereas LMNC levels did not differ by genotype. On further examination, we detected additional species with the LMNA/C antibody (55kD) indicating most likely cleavage of LMNC in the STIM1^+/D84G^ nuclear extracts (26) (Figure 6A). These LMNC fragments migrated as high molecular weight multimers under non-reducing PAGE conditions (85-150 kD). In addition, this 55 kD LMNA/C fragment was detected to a greater extent in the soluble cytosolic fraction of the STIM1^+/D84G^ muscles, findings consistent with LMNC damage (Supplemental Data Figure 1A). To further test whether STIM1 D84G disrupted LMNA/C and cause nuclear export of LMNA/C, we compared LMNA-GFP localization in HEK-293 cells expressing with either the STIM1 WT or the STIM1 D84G. LMNA-GFP was detected outside of the nucleus to a greater extent for cells expressing STIM1 D84G (Supplemental Data Figure 1C). We interpret these data as evidence that the D84G mutant STIM1 in NE destabilized the nuclear lamina and LMNA/C filamentous network (Supplemental Figure 2A).

**Figure 6:**
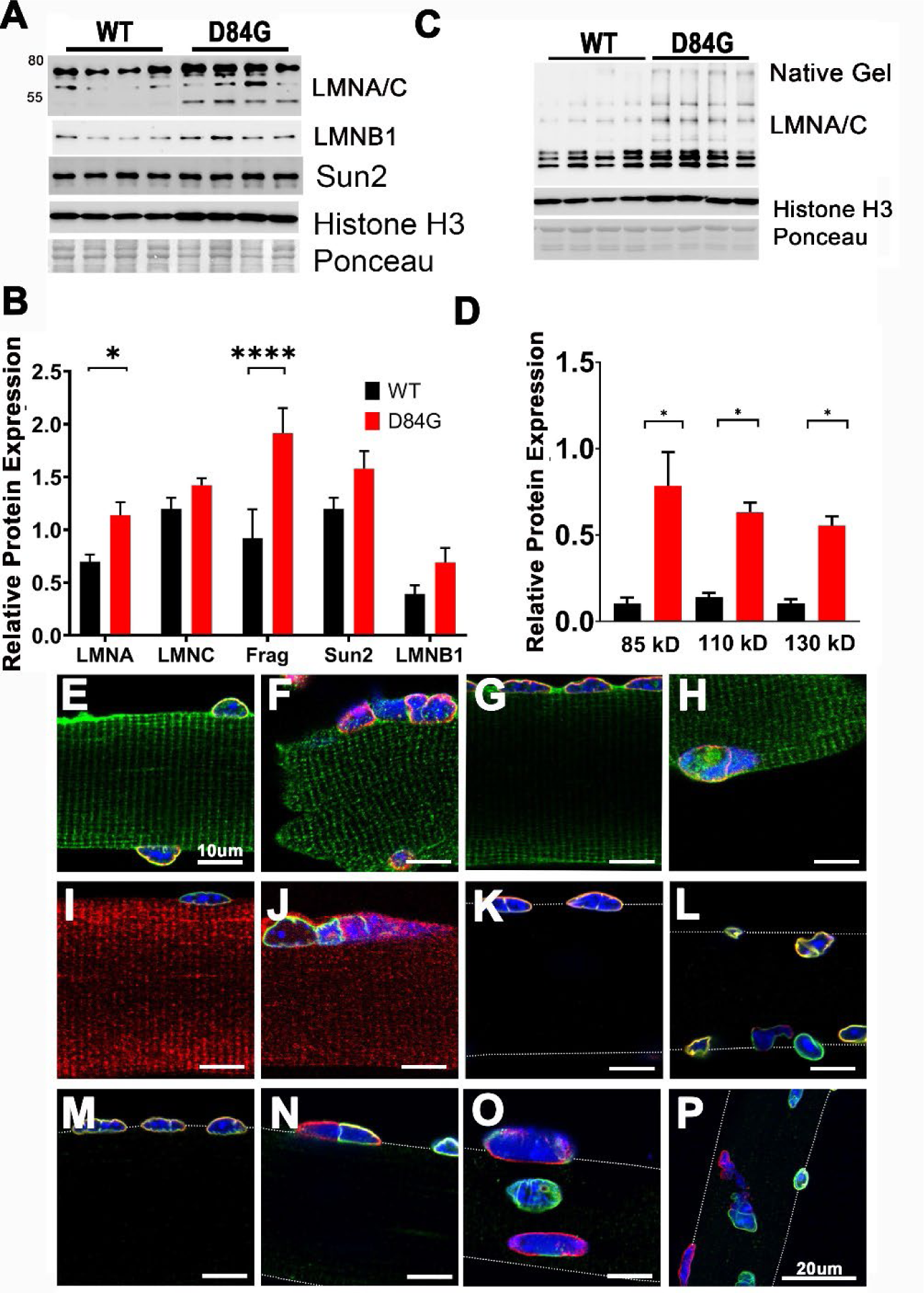
Nuclear membrane proteins in WT and D84G mice. Nuclear extracts from gastrocnemius muscle were subjected to SDS-PAGE and immunoblotting for WT (n=4) and STIM1^+/D84G^ mice (n=4). (A) Immunoblotting for LMNA/C, LMNB1, Sun2 and Histone H3. LMNA/C blots is represented by isoforms (60-80 kD) and smaller fragments (55 kD). (C) Immunoblotting for LMNA/C under non-reducing conditions by removing DTT demonstrate aggregates of LMNA fragments (85, 110 and 130 kD). (B-D) Quantification of the A) and C) of proteins in WT (black) and STIM1^+/D84G^ (gray) muscles. Values are mean ± standard deviation. (E-P) Localization of nuclear membrane proteins in FDB fibers by fluorescence immunohistochemistry (n=3-5 mice). (E-F) STIM1 (green) LMNA/C (red) in WT (E) and STIM1^+/D84G^ mice (F). (G-H) STIM1 (green) SUN2 (Red) in WT (G) and STIM1^+/D84G^ mice (H). (I-J) LMNB1 (green) and STIM1 (red) in WT (I) and STIM1^+/D84G^ mice (J). (K-L) LMNB1 (Green)SUN2 (Red) in (K) WT and (L) STIM1^+/D84G^ mice. (M-P) LMNB1 (green) LMNA/C (Red) in (M) WT and (N-P) STIM1^+/D84G^ mice. (E-O) are confocal images taken at 40x with a 5x zoom, scale bars 10um. (P) is a maximum image projection of 11 confocal images taken at 2um intervals through the fiber at 40x with 3x zoom, scale bars 20um. Two-tailed Student’s *t*-test, n=3-5 independent experiments. *p<0.01

Co-staining FDB fibers for STIM1 along with LMNA/C (red)(Figure 6E-F), SUN2 (red) (Figure 6 G-H) or LMNB1 (green) (Figure 6I-J) demonstrate co-localization of the LMNA/C, LMNB1 and SUN2 with STIM1 in WT nuclear membrane. In the majority of nuclei in STIM1^+/D84G^ fibers STIM1 also co-localizes with LMNA/C, SUN2 and LMNB1. However, there are numerous nuclei with marked disorganization of the nuclear lamina in STIM1^+/D84G^ fibers. Lamin A/C (red) staining was found to be absent in a number of dysmorphic nuclei (Figure 6E-F). SUN2 (red) was generally present in all nuclear membranes but downregulated in some nuclei with disrupted morphology (Figure 6G-H). LMNB1 (green) was, again, not present in all nuclei of STIM1^+/D84G^ muscles and was notably absent from micronuclei (Figure 6I-J, 6P). To better understand the differences to the nuclear lamina in the D84G fibers, we co-stained for LMNA/C (red) and SUN2 (red) with LMNB1 (green). In contrast to WT nuclei, where there was good co-localization for LMNB1 and SUN2 in muscle nuclei, LMNB1 (green) staining was often lost in portions of misshapen nuclei from the STIM1^+/D84G^ fibers, SUN2 was evident in all nuclei but was down regulated in some regions of NE from STIM1^+/D84G^ nuclei. (Figure 6 K-L). Similarly, LMNA/C (red) was often detected independent of the LMNB1 (green) in nuclei of STIM1^+/D84G^ fibers, but exhibited good colocalization in WT nuclei (Figure 6 M-P). These data are consistent with the notion that nuclear lamina was damaged in nuclei expressing the D84G mutant STIM1 as evident by the altered spatial localization for nuclear lamina and the LINC complex.

### LMNA-dependent gene expression is altered in STIM1^+/D84G^ muscle

To this point, our findings show that the nuclei of STIM1^+/D84G^ muscle are distorted and nuclear lamina damaged, features that resemble NE subjected to excessive mechanical forces or as part of the pathology seen in laminopathies. We hypothesized that changes to the nuclear lamina of the STIM1^+/D84G^ muscle would interrupt LMNA-chromatin interactions leading to expression of genes that are otherwise silenced, as has been described in LMNA KO muscles (27, 28). Consistent with this idea, GSEA analysis of our RNA-sequence data from STIM1^+/D84G^ muscles revealed negative enrichment of genes regulating DNA repair and chromatin modification (Figure 7A). To confirm this idea, we quantified a set of genes known to be upregulated in the LMNA KO muscles and found profound changes in these mRNAs were detected in the muscles of STIM1^+/D84G^ mice including for sarcolipin (40-fold) (SLN), Ankryin domain repeat 1 (18-fold) (ANKRD1), microsomal triglyceride transfer protein (2-fold) (MTTP), Ca^2+^ and integrin binding proteins (0.5 fold reduction) (CIB2) and S100 Ca^2+^ binding protein A4 (9-fold increase) (S100A4)(Figure 7B). Collectively these data indicate that D84G STIM1 in the NE likely leads to an alteration in the lamina-associated chromatin structure which may account for the large number of DEGs in the STIM1^+/D84G^ muscles.

**Figure 7:**
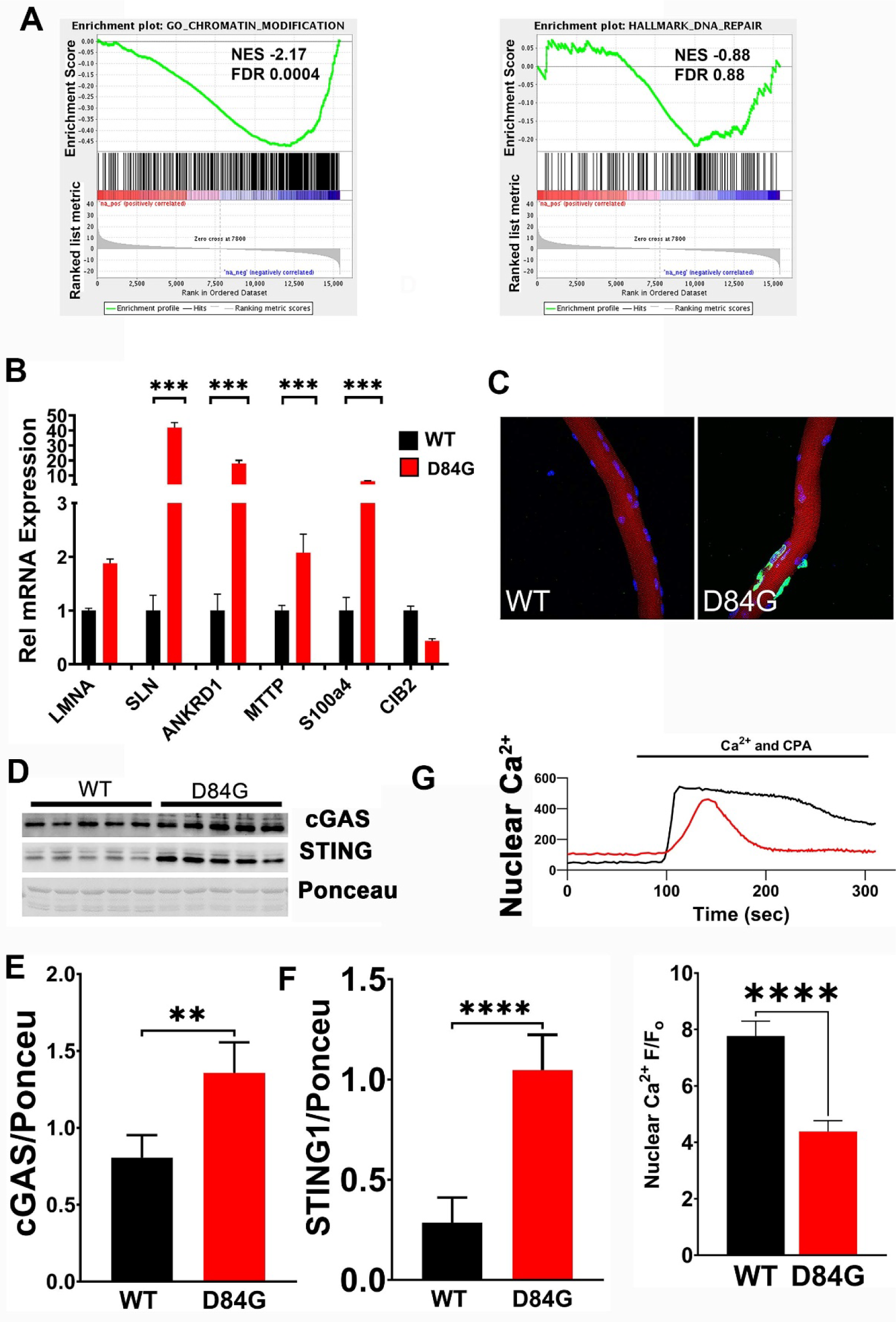
Muscle fibers from STIM1^+/D84G^ mice exhibit nuclear dysfunction. (A) GSEA WT vs. STIM1^+/D84G^ RNA-Seq. FDR q value and P value are shown for each plot. Negative enrichment of gene sets involved DNA repair and chromatin organization are shown. (B) LMNA-target gene expression in the WT and STIM1^+/D84G^ muscle. We mined our RNA-seq data (from WT and STIM1^+/D84G^ muscle) for genes known to be differentially expressed in LMNA KO muscle, as described in (27, 28). Quantification of results (Mean ± S.E.) represents n=4 animals for each genotype. *** P<0.01 by Student’s t-test significant only for STIM1^+/D84G^ versus WT. (C) FBD fibers for WT and D84G mice (C) were fixed and immunostained with the γ-H2A.X antibody (green) to detect phosphorylated Histone H2A levels, a marker of DNA damage. Nuclei are co-labeled with DAPI (blue). Contours of the fiber were demonstrated by phalloidin to label actin (red). Results are representative for three mice of each genotype. (D-F) Nuclear lysates from 6-month mice were prepared from WT (n=5) and STIM1^+/D84G^ (n=5) muscles and immunoblotted for cGAS and STING with specific antibodies. Ponceau staining were used for loading control. Quantification of results (Mean ± S.E.) represents n=5 animals for each genotype. ** P<0.01, **** P<0.001 by Student’s t-test significance for STIM1^+/D84G^ versus WT. (G) WT and D84G STIM1 myoblasts were transfected with gCAMP genetically encoded Ca^2+^ indicator. SERCA inhibitor cyclopiazonic acid (30uM) was used to evoke Ca^2+^ transients and then Ca^2+^ was added back to refill [Ca^2+^]_N_. Comparison of the nuclear Ca^2+^ content was determined as the F/F_0_. Results are representative of n=50 cells for WT and D84G mutants each. Imaging was performed on 5 separate days. Results are Mean ± S.E.**P<0.01, ****P<0.001 by Student’s t-test significant for STIM1^+/D84G^ versus WT STIM1 transfected myoblasts.

### DNA damage in the STIM1^+/D84G^ muscles

The foregoing observations that muscle fibers expressing the STIM1 D84G mutant exhibit damage to nuclear lamina and altered LMNA/C gene expression raised the possibility that DNA damage may accumulate in the nuclei of STIM1^+/D84G^ muscle and thereby represent an important mechanism underlying reduced muscle growth and weakness. Genomic instability can be detected at sites of double stranded breaks in DNA in muscle nuclei as an increase in phosphorylation of Histone H2A.X. DNA damage was detected by the presence of γH2A.X positive (γH2AX+) myonuclei in STIM1^+/D84G^ muscle fibers. While no γH2AX+ fibers were detected in the WT muscles, myonuclei from STIM1^+/D84G^ muscle exhibited significant number of γH2A.X+ myonuclei indicating D84G STIM1 associates with nuclear lamina damage and cause DNA damage (Figure 7C). The γH2AX+ myonuclei were most often the misshapen nuclei, consistent with loss of the integrity of NE. Additional evidence for DNA damage in the D84G mutant fibers included demonstration of the upregulation for the enzyme cyclic GMP-AMP synthase (cGAS) which is a DNA sensor (Figure 7D-E). Similarly, western blotting for stimulator of interferon genes (STING1) was increased in muscles of STIM1^+/D84G^ mice (Figure 7D and F). STING is activated by cytosolic DNA and cGAS where it contributes to DNA damage signaling and activation of the sterile inflammation cascade. Together these data show that presence of D84G STIM1 in the NE creates proteostatic stress and DNA damage which impairs muscle growth and performance as seen in the STIM1^+/D84G^ mice.

### D84G STIM1 reduced nuclear Ca^2+^

We next wanted to know if the D84G mutation in STIM1 influenced [Ca^2+^]_N_. In resting muscle cells, [Ca^2+^]_N_ is maintained by tonic leak of Ca^2+^ from IP3R channels and Ca^2+^ efflux by the SERCA1(29-31). In contrast, the nuclear pores (NPC) act as passive conduits for Ca^2+^ during cell stimulation and determine of the amplitude of [Ca^2+^]_N_(32, 33). The genetically encoded Ca^2+^ sensor gCAMP6f can be targeted to the nucleus by the addition of a SV40 nuclear localization sequence(34). We expressed n-gCAMP6 in myoblasts expressing either WT or D84G STIM1. Myoblasts were subjected to Ca^2+^ store depletion by application of the SERCA inhibitor CPA in the absence of external Ca^2+^, a protocol known to activate STIM1. Upon readdition of Ca^2+^, [Ca^2+^]_N_ increased for both WT and STIM1 D84G myoblasts, as Ca^2+^ enter the nucleus by the actions of the NPC. Under these conditions, the amplitude of [Ca^2+^]_N_ was significantly reduced in the D84G-STIM1 expressing muscle cells compared WT (Figure 7G). These findings are consistent with the idea that D84G STIM1 influenced Ca^2+^ flux across the NE, most likely mediated by the NPC. Consistent with this idea, we found significant reduction in the mRNA transcript reads for several components of the NPC in the RNA seq data set from STIM1^+/D84G^ muscles (Supplemental Figure 2B).

## Discussion

In the present work, we provide compelling evidence that a gain of function mutation in the STIM1 gene is sufficient to induce a progressive myopathy accompanied by minor changes in Ca^2+^ signaling and drastic changes to nuclear structure and function. Specifically, we show that mice harboring a single nucleotide change in STIM1 (D84G) display (i) constitutive Ca^2+^ entry across the sarcolemma of muscle fibers, (ii) reduced muscle mass and weakness consistent with sarcopenia, (iii) and disruption of NE resulting in altered nuclear architecture, gene expression and DNA-damage. Overall, these studies establish that STIM1^+/D84G^ mice replicate the clinical aspects of weakness and reduced muscle mass that occur in humans and offers new insight to the function of STIM1 in NE and highlight how STIM1 links mechanical signals with gene expression, DNA damage and muscle growth.

In skeletal muscle, STIM1 localizes to specialized SR domains where it interacts with different target proteins to carry out distinct cellular functions. For example, STIM1 is present in the membrane of the SR terminal cisternae where it activates Orai1 channels in the adjacent T-tubule membrane (4). Far less is known about the role of STIM1 in the NE of muscle cells (12). We detected STIM1 in the NE and throughout the invaginations of the NE called the nuclear reticulum, as was previously shown (35). In an attempt to understand the function of STIM1 has in NE, we characterized the nuclear phenotype of the STIM1^+/D84G^ mice. TEM of muscles from STIM1^+/D84G^ mice demonstrate expansion of the space between the ONM and INM (perinuclear space, PNS) as well as altered nuclear morphology. These findings are reminiscent of prior studies from cells depleted of SUN proteins that exhibit enlarged PNS due to loss of key connections between SUN1 and SYNE1 in the ONM (36). In SUN1/SUN2 KO mice, defects in myonuclei anchorage results from the failure to retain Nesprin-1 at the ONM. The mutant STIM1 present in the ONM may also disrupt key components of the LINC complex and thereby activate nuclear stress signaling. Whether STIM1 connects the NE to the cytoskeleton where it may integrate mechanical signaling requires further investigation. In fact, we recently demonstrated that STIM1 interacts with Desmin, a muscle specific cytosolic intermediate filament. Given that Desmin is known to interact with SYNE1 to stabilize nuclear positioning and maintain nuclear architecture of muscle, we propose that STIM1 and Desmin establish another important SR domain in muscle, in this case with the NE where it is involved in mechanotransduction for the muscle (37, 38).

Disruption of the folding of the EF hand for STIM1 by the D84G mutation eliminates the Ca^2+^ sensor function of STIM1 leading to constitutive SOCE in muscle fibers. Importantly, muscle fibers adapt to the constitutive SOCE as there was no change to Ca^2+^ stores, resting Ca^2+^ or EFS-evoked Ca^2+^ transients. The upregulation of Sarcolipin and Calsequestrin in STIM1^+/D84G^ muscle provide adaptation to the augmented SOCE by enhancing Ca^2+^ storage and limiting SERCA1 refilling. In contrast, our studies show that cytosolic transfer of Ca^2+^ into the nucleus by NPC was reduced when D84G STIM1 was expressed myoblasts (39, 40). The consequence of the lower [Ca^2+^]_N_ in the STIM1^+/D84G^ myoblasts is likely due to altered mechanical properties of the nucleus. These data are consistent with recent studies where mechanical stress disrupted the link between LMNA/C, lamina associated chromatin and nuclear Ca^2+^ levels that subsequently influence chromatin structure, chromatin accessibility and gene expression (41). Our RNA sequencing data from muscles of 6-month old STIM1^+/D84G^ mice demonstrated a very significant ∼10% increase in differentially expressed genes. These changes likely reflect disruption of chromatin contacts with the lamina as was reflected by the GO analysis and the enhanced expression of LMNA/C regulated genes. Nuclear periphery is often aligned with condensed chromatin that harbors genes that are transcriptionally silent(42). We therefore interpret our studies as evidence that nuclei of STIM1^+/D84G^ muscle resemble nuclei in WT muscle subjected to excessive stretch where heterochromatin structure is modified, and ER/nuclear Ca^2+^ altered resulting in DNA damage (43). This work therefore establishes a previously unappreciated role for STIM1 in regulating NE dynamics and possible role in mechanotransduction and gene expression.

The clinical syndrome of muscle atrophy and weakness develops in humans as a result of gain of function mutations in the STIM1 and Orai1 genes. Pathologic biopsy specimens and MRI imaging demonstrate TAs in muscles of these patients has led to the assertion that constitutive SOCE causes TA aggregates (44). Studies to date fail to demonstrate TAs in the muscles of STIM1 and Orai1 mouse models despite the presence of constitutive SOCE, reduced muscle growth and muscle weakness (5-9, 16, 44-52). In contrast, TAs are often detected in muscles of aging WT mice where the regulation of SOCE is likely altered. While upregulation of SOCE has been proposed to occur in aging muscle, it remains to be determined if STIM1-dependent Ca^2+^ signaling is coincidently enhanced in the aging muscle (53, 54). Results of these studies question the relationship of STIM1 and Orai1 mutations as the cause of TAs. Based on our work with this STIM1 mutant mouse model, we propose STIM1 senses mechanical stress as well as Ca^2+^ stores content in the NE and nuclear lamina. STIM1 can then influence nuclear Ca^2+^ flux through the NPC, limit LMNA-chromatin changes and prevent DNA damage. This work establishes a previously unrecognized role for STIM1 signaling in the nuclear dynamics. Finally, it is important to note the relevance of this signaling pathway to aging and that STIM1 has a physiologic role in sinus node pacemaking and arrhythmias, immune responsiveness and muscle metabolism; all of which are perturbed with aging. The implication that STIM1 is a potential therapeutic target for aging would therefore require further study.

## Materials and Methods

### Animals

*Stim1^+/D84G^* mice were generated as described^7^. Sequencing of this mouse line shows an A-to-G transition at nucleotide 444 in exon 2 of *Stim1* (NM_009287), resulting in an amino acid exchange in the EF hand motif at position 84 (Asp84Gly, or D84G). Age and gender matched C57BL/6 mice were used as control wild types (WT). All mice were maintained in pathogen-free barrier facilities at Duke University and were used in accordance with protocols approved by the Division of Laboratory Animal Resources and Institutional Animal Care & Use Committee at Duke University.

### Isolation of FDB fibers

FDB muscles were carefully dissected from the foot and placed into DMEM with 0.5-1mg/ml Collagenase A (Roche 10103578001). Fibers were digested at 37°C overnight, with rocking, before isolation by gentle trituration in DMEM using a glass pipette. Once isolated, fibers were plated on 35mm glass bottomed dishes coated with 20ug/ml laminin (Sigma - L2020).

### Nuclear Enrichment

Frozen skeletal muscle tissue was powdered with liquid nitrogen and re-suspended in SMT buffer (0.2% NP40, 250mM Sucrose, 50mM Tris.cl, 5mM MgCl2 pH 7.6 with protease inhibitors) for homogenization and centrifuged at 1,000g and the pellet were collected. The nuclear pellet was washed by SMT buffer three times and resuspended in NET buffer (20mM Tris.Cl, 150mM NaCl, 1.5mM MgCl2,1% SDS and 5% glycerol, pH 7.6 with protease inhibitor), vortexed 15sec and passed through a 20-gauge needle and then centrifuged at 11,000g for 15min to collect the nuclear lysate.

### Immunoblotting

Standard protocols were used for immunoblot analysis. Blots were incubated overnight at 4°C with primary antibodies. Antibodies against STIM1 (Sigma cat#6197) SERCA1a (Thermofisher, ma3912 mouse), RyR (Thermofisher, 3925), Casquestrin (Thermodisher, 3913 mouse), SLN (Millipore, ABT13), LaminA/C (Cell signaling antibody 4C11, Mouse mAb), SUN2 (Millipore MABT880), cGAS (Cell signaling, 31695), STING1 (Cell signaling, 50494), Histone H3 (Cell signaling, 9715s) and GAPDH (Sigma, G9545). Signals were detected using horseradish peroxidase (HRP)-conjugated secondary antibody and Pierce super signal western Dual extended duration substrate kit (34075), and quantified with the ChemiDoc™MP Imaging System (Bio-Rad).

## Histology

Fibers were fixed in 4% PFA for 5 minutes, washed in PBS and blocked in 10% heat inactivated goats serum (HINGS). Fibers were incubated with primary antibody overnight at room temperature in fiber antibody solution (PBS, 2% HINGS and 0.3% Triton). Primary antibodies are outlined in the supplemental table 2. After washing, fibers were incubated with secondary antibody for 1h (molecular probes – Alexa Fluor series), washed, and mounted in VECTASHIELD. Nuclei in whole fibers were stained for 5 minutes with 10ug/ml DAPI. Staining was analyzed on a Zeiss 510 confocal microscope. For staining of cryosections on slides, a similar protocol was used, but the antibody solution was 1% HINGS with 0.1% Triton.

## Electron Microscopy

For ultrastructural localization of STIM1-LacZ by TEM, FDB muscles were fixed in situ for 5 minutes in 2% PFA 0.2% Gluteraldehyde in PBS. Nodes were then dissected and stained for LacZ as described in Stiber et al. (Stiber et al., 2008). Tissue was post fixed in 2% PFA and 2% glutaraldehyde in 0.1 M phosphate buffer, pH 7.4. then fixed in 1% osmium tetraoxide, and stained en bloc with 1% uranyl acetate. Tissue was then dehydrated in a graded ethanol series, taken through a series of Spurr resin/ethanol washes and embedded in Spurr resin. Thin sections were cut at 70nm, mounted on copper grids and counterstained with 2% uranyl acetate and lead citrate. Grids were viewed and photographed using a FEI Tecnai G2 Twin transmission electron microscope. Semi-thin sections were cut at 1um and mounted on slides and dried on a hotplate. They were stained with 1% Toluidine Blue and 2% Borate in distilled water with the hotplate still on. Excess stain was removed with distilled water and dried. Slides were cover slipped in Cytoseal 60.

## Calcium Imaging of FDB Myofibers

### Manganese quench

Fibers were loaded with 1ug/ml FURA-2AM for 30 minutes in Tyrode’s solution (121 mM NaCl, 5mM KCl, 1.8mM CaCl2, 500μM MgCl2, 400μM NaH2PO4, 100μM EDTA, 5.5mM glucose, and 24mM NaHCO3). Solutions were bubbled with oxygen throughout. FURA-2AM was removed by perfusing with Tyrode’s solution with 50uM BTS, to prevent motion artifact. During this period of time healthy fibers were selected by applying a single stimulus for 100ms (100 mA) at 50 Hz. Fibers were imaged for 3 minutes in Tyrode’s solution with Ca2+ to determine the basal level of FURA-2. The Ca2+ was subsequently replaced with 1.8 mM Mn by perfusion for 5 minutes at 1ml/minute. Fibers were then stimulated with 20 trains at 50 Hz, 1s duration every 5 seconds. 360nm using an Acupulser 310 (World Precision Instruments) with an A385 Stimulus Isolator (World Precision Instruments). Stimulation dish was purchased from World Precision Instruments RC-37FS. Florescence signal was monitored throughout the experiment using an excitation filter at 360, emission filter 510. Images were collected every 2 seconds on a Nikon TE2000 inverted microscope equipped with a Photometrics CoolSNAP camera and a Lambda DG-4 rapid filter changer (Sutter). Image acquisition was controlled using Metafluor/Metamorph software (Molecular Devices). Images were acquired using a 40× S Plan 1.3-numerical-aperture (NA) objective lens.

### Basal Calcium

Fibers were loaded with 1ug/ml FURA-2AM for 30 minutes in oxygenated imaging solution (120mM NaCl, 5mM KCl, 2mM CaCl2, 1mM MgCl2, 25 mM NaHCO3, 0.5mM NaH2PO4 and 10mM glucose). FURA-2 was washed out of the dish for 5 minutes and fibers selected for viability. Ca2+ signals were measured by alternate excitement at 340nm and 380nm emission at 510nm. Acquisition was every 1s on a PCO Edge 5.5 camera, core LED 340 FURA light source. Image acquisition was controlled using Metafluor/Metamorph software (Molecular Devices). Images were acquired using a 40× S Plan 1.3-numerical-aperture (NA) objective lens.

### Fiber Stimulation

Fibers were loaded with 2ug/ml FURA-4F AM for 1h in oxygenated imaging solution. FURA-4F AM was removed by perfusion with imaging solution with BTS for 5 minutes. Fibers were selected for viability then subjected to a stimulation protocol whilst being perfused with oxygenated imaging solution. A stimulus response curve was generated by applying stimuli ranging from 100ms -2s every 45 seconds. After this, trains of stimuli were applied with 25 stimuli of 500ms every 5 seconds first then 25 stimuli of 2s every 5 seconds. Ca2+ signals were measured by alternate excitement at 340nm and 380nm emission at 510nm. Acquisition was every 180ms 340nm and 80ms 380nm on a PCO Edge 5.5 camera, core LED 340 FURA light source). Camera and software was as described for basal calcium measurements.

### Store Depletion

Fibers were loaded with 2ug/ml FURA-4F AM for 1h in oxygenated imaging solution. FURA-4F AM was removed by perfusion with imaging solution with BTS for 5 minutes. Fibers were selected for viability and basal calcium recorded. Fibers were then perfused with imaging solution with 0 calcium for 3 minutes. Stores were depleted using a solution with 0 calcium, 30um CPA and 10 mm caffeine.

### Nuclear Ca^2+^ Measurements

To measure nuclear Ca^2+^, plasmids encoding WT and D84G STIM1 were transfected to C2C12 cells with the genetically encoded Ca^2+^ indicator pEGFP-N1-GCaMP6m-Xn (a gift from Xiaodong Liu, Addgene plasmid # 118976; http://n2t.net/addgene:118976; RRID:Addgene_118976) targeted to the nucleus. Wide field epifluorescence was recorded for the GFP signal. Myoblasts were perfused with solutions HBSS (Gibco 14175-095) with 1mM Mg^2+^ for the SOCE assay including no Ca^2+^ solution, no Ca^2+^ with cyclopiazonic acid (30uM) to deplete stores and then add back 2 mM Ca^2+^ with CPA. Change in nuclear Ca^2+^ was determined from the peak of signal after Ca^2+^ add back normalized to the average lowest level in no Ca^2+^ CPA.

## Behavioral Studies

**Open field activity** was monitored using the VersaMax Animal Activity Monitoring System as previously described33. Mice were tested in individual chambers for a total of 15 minutes before and immediately after exercise. Every 5 minutes of ambulation, the average total distance and time spent in movement as well as vertical activity was determined and standard errors were calculated. Cumulative measures for every 5 minute interval were charted. Mice were exercised with an adjustable variable-speed belt treadmill from AccuPacer as previously described33.

### Grip strength

Measurements were done for each individual mouse using a San Diego Instruments animal grip strength system. Grip strength readings Maximum force (N) were taken in triplicate and hind paw strength was calculated indirectly after measurements of front paw and whole-body strength.

### Involuntary running

Mice were trained to run on the treadmill for 3 days at a speed of 5 meters/minute for 10 minutes. When they stopped running, they were tapped lightly to encourage them to restart. For the experimental run, mice started at a speed of 5 meters/minute for 2 minutes, the speed was increased by 1 meter/minute until the mice were no longer able to run. The end point was taken when mice were tapped 3 times and failed to restart.

## Real-time PCR Analysis and RNA-seq

RNA was isolated using a TRIzol extraction protocol followed by clean up using an RNeasy Mini Kit. Total mRNA was reversed transcribed into cDNA using the Applied Biosystems High-Capacity mRNA to cDNA kit. cDNA was amplified using TaqMan Gene Expression Master Mix and TaqMan Gene Expression Assays. Taqman Primers from Thermo used were as follows: BIP/HSPA5 mm00517691, PIDA4 mm00437958, SDF21L mm00452079, GADD45 45 mm00432079, Myogennin mm00446194, Pax7 mm01354484, Myostatin mm01254459, CrystAB mm00515567, GAPDH mm99999915, 18S Hs99999901.

The Sequencing and Genomics Technologies Core Facility at Duke University performed mRNA-sequencing. The SGT used the Kapa stranded mRNA Kit from Roche (Code: KK8421) to enrich for mRNA from total RNA and reversed transcribed mRNA into cDNA to build sequencing libraries. Libraries were pooled to equimolar concentration and sequenced on the NovaSeq 6000 SP flow cell to produce 50 bp paired-end reads.

### Statistical Analysis

The numbers of mice used in experiments for each study are indicated in figure legends. Values are presented as means ± standard errors of the means (SEM). A two-tailed Student’s *t* test was used to calculate *P* values (significant if <0.05).

### Data Availability

The data that support this study are available from the corresponding author upon reasonable request. Raw RNA-sequencing data generated in this study have been deposited in the Gene Expression Omnibus (GEO) under accession number XXXXX.

## Supporting information

all suppl figures

## Acknowledgements

We thank the Duke Core for Electron Microscopy and the Duke Center for Genomic and Computational Biology with technical assistance. This work was supported by the funding from the National Institute of Health 5R01-DK109911 and 5R01-HD096385 (PBR).

## Notes

### Competing Interest Statement

The authors have declared no competing interest.

## References

1. Rosenberg PB. Calcium entry in skeletal muscle. J Physiol. 2009;587(Pt 13):3149–51.

2. Lyfenko AD, and Dirksen RT. Differential dependence of store-operated and excitation-coupled Ca2+ entry in skeletal muscle on STIM1 and Orai1. J Physiol. 2008;586(Pt 20):4815–24.

3. Wei-Lapierre L, Carrell EM, Boncompagni S, Protasi F, and Dirksen RT. Orai1-dependent calcium entry promotes skeletal muscle growth and limits fatigue. Nat Commun. 2013;4:2805.

4. Stiber J, Hawkins A, Zhang ZS, Wang S, Burch J, Graham V, et al. STIM1 signalling controls store-operated calcium entry required for development and contractile function in skeletal muscle. Nat Cell Biol. 2008;10(6):688–97.

5. Bohm J, Chevessier F, De Paula AM, Koch C, Attarian S, Feger C, et al. Constitutive Activation of the Calcium Sensor STIM1 Causes Tubular-Aggregate Myopathy. Am J Hum Genet. 2013.

6. Noury JB, Bohm J, Peche GA, Guyant-Marechal L, Bedat-Millet AL, Chiche L, et al. Tubular aggregate myopathy with features of Stormorken disease due to a new STIM1 mutation. Neuromuscul Disord. 2017;27(1):78–82.

7. Harris E, Burki U, Marini-Bettolo C, Neri M, Scotton C, Hudson J, et al. Complex phenotypes associated with STIM1 mutations in both coiled coil and EF-hand domains. Neuromuscul Disord. 2017;27(9):861–72.

8. Bohm J, Bulla M, Urquhart JE, Malfatti E, Williams SG, O’Sullivan J, et al. ORAI1 Mutations with Distinct Channel Gating Defects in Tubular Aggregate Myopathy. Hum Mutat. 2017.

9. Okuma H, Saito F, Mitsui J, Hara Y, Hatanaka Y, Ikeda M, et al. Tubular aggregate myopathy caused by a novel mutation in the cytoplasmic domain of STIM1. Neurol Genet. 2016;2(1):e50.

10. Lee JM, and Noguchi S. Calcium Dyshomeostasis in Tubular Aggregate Myopathy. Int J Mol Sci. 2016;17(11).

11. Zhang H, Bryson VG, Wang C, Li T, Kerr JP, Wilson R, et al. Desmin interacts with STIM1 and coordinates Ca2+ signaling in skeletal muscle. JCI Insight. 2021;6(17).

12. Lee SH, Hadipour-Lakmehsari S, Miyake T, and Gramolini AO. Three-dimensional imaging reveals endo(sarco)plasmic reticulum-containing invaginations within the nucleoplasm of muscle. Am J Physiol Cell Physiol. 2018;314(3):C257–C67.

13. Iyer SR, Folker ES, and Lovering RM. The Nucleoskeleton: Crossroad of Mechanotransduction in Skeletal Muscle. Front Physiol. 2021;12:724010.

14. Madej-Pilarczyk A. Clinical aspects of Emery-Dreifuss muscular dystrophy. Nucleus. 2018;9(1):268–74.

15. Grosse J, Braun A, Varga-Szabo D, Beyersdorf N, Schneider B, Zeitlmann L, et al. An EF hand mutation in Stim1 causes premature platelet activation and bleeding in mice. J Clin Invest. 2007;117(11):3540–50.

16. Bohm J, Chevessier F, Koch C, Peche GA, Mora M, Morandi L, et al. Clinical, histological and genetic characterisation of patients with tubular aggregate myopathy caused by mutations in STIM1. J Med Genet. 2014;51(12):824–33.

17. Boncompagni S, Protasi F, and Franzini-Armstrong C. Sequential stages in the age-dependent gradual formation and accumulation of tubular aggregates in fast twitch muscle fibers: SERCA and calsequestrin involvement. Age (Dordr). 2012;34(1):27–41.

18. Zhang SL, Yu Y, Roos J, Kozak JA, Deerinck TJ, Ellisman MH, et al. STIM1 is a Ca2+ sensor that activates CRAC channels and migrates from the Ca2+ store to the plasma membrane. Nature. 2005;437(7060):902-5.

19. Stathopulos PB, Zheng L, Li GY, Plevin MJ, and Ikura M. Structural and mechanistic insights into STIM1-mediated initiation of store-operated calcium entry. Cell. 2008;135(1):110–22.

20. Koenig X, Choi RH, and Launikonis BS. Store-operated Ca(2+) entry is activated by every action potential in skeletal muscle. Commun Biol. 2018;1:31.

21. Li T, Finch EA, Graham V, Zhang ZS, Ding JD, Burch J, et al. STIM1-Ca(2+) signaling is required for the hypertrophic growth of skeletal muscle in mice. Mol Cell Biol. 2012;32(15):3009–17.

22. Odermatt A, Becker S, Khanna VK, Kurzydlowski K, Leisner E, Pette D, et al. Sarcolipin regulates the activity of SERCA1, the fast-twitch skeletal muscle sarcoplasmic reticulum Ca2+-ATPase. J Biol Chem. 1998;273(20):12360–9.

23. Huang da W, Sherman BT, and Lempicki RA. Systematic and integrative analysis of large gene lists using DAVID bioinformatics resources. Nature protocols. 2009;4(1):44–57.

24. Sherman BT, Hao M, Qiu J, Jiao X, Baseler MW, Lane HC, et al. DAVID: a web server for functional enrichment analysis and functional annotation of gene lists (2021 update). Nucleic Acids Res. 2022;50(W1):W216-21.

25. Mattout A, Dechat T, Adam SA, Goldman RD, and Gruenbaum Y. Nuclear lamins, diseases and aging. Curr Opin Cell Biol. 2006;18(3):335–41.

26. Takahashi A, Alnemri ES, Lazebnik YA, Fernandes-Alnemri T, Litwack G, Moir RD, et al. Cleavage of lamin A by Mch2 alpha but not CPP32: multiple interleukin 1 beta-converting enzyme-related proteases with distinct substrate recognition properties are active in apoptosis. Proc Natl Acad Sci U S A. 1996;93(16):8395–400.

27. Wang WP, Wang JY, Lin WH, Kao CH, Hung MC, Teng YC, et al. Progerin in muscle leads to thermogenic and metabolic defects via impaired calcium homeostasis. Aging Cell. 2020;19(2):e13090.

28. Chandran S, Suggs JA, Wang BJ, Han A, Bhide S, Cryderman DE, et al. Suppression of myopathic lamin mutations by muscle-specific activation of AMPK and modulation of downstream signaling. Hum Mol Genet. 2019;28(3):351–71.

29. Zima AV, Bare DJ, Mignery GA, and Blatter LA. IP3-dependent nuclear Ca2+ signalling in the mammalian heart. J Physiol. 2007;584(Pt 2):601–11.

30. Stiber JA, Tabatabaei N, Hawkins AF, Hawke T, Worley PF, Williams RS, et al. Homer modulates NFAT-dependent signaling during muscle differentiation. Dev Biol. 2005;287(2):213–24.

31. Molgo J, Colasantei C, Adams DS, and Jaimovich E. IP3 receptors and Ca2+ signals in adult skeletal muscle satellite cells in situ. Biological research. 2004;37(4):635–9.

32. Malhas A, Goulbourne C, and Vaux DJ. The nucleoplasmic reticulum: form and function. Trends Cell Biol. 2011;21(6):362–73.

33. Bootman MD, Fearnley C, Smyrnias I, MacDonald F, and Roderick HL. An update on nuclear calcium signalling. J Cell Sci. 2009;122(Pt 14):2337–50.

34. Enyedi B, Jelcic M, and Niethammer P. The Cell Nucleus Serves as a Mechanotransducer of Tissue Damage-Induced Inflammation. Cell. 2016;165(5):1160–70.

35. Vahabikashi A, Sivagurunathan S, Nicdao FAS, Han YL, Park CY, Kittisopikul M, et al. Nuclear lamin isoforms differentially contribute to LINC complex-dependent nucleocytoskeletal coupling and whole-cell mechanics. Proc Natl Acad Sci U S A. 2022;119(17):e2121816119.

36. Crisp M, Liu Q, Roux K, Rattner JB, Shanahan C, Burke B, et al. Coupling of the nucleus and cytoplasm: role of the LINC complex. J Cell Biol. 2006;172(1):41–53.

37. Heffler J, Shah PP, Robison P, Phyo S, Veliz K, Uchida K, et al. A Balance Between Intermediate Filaments and Microtubules Maintains Nuclear Architecture in the Cardiomyocyte. Circ Res. 2020;126(3):e10–e26.

38. Rey A, Schaeffer L, Durand B, and Morel V. Drosophila Nesprin-1 Isoforms Differentially Contribute to Muscle Function. Cells. 2021;10(11).

39. Magli E, Fattorusso C, Persico M, Corvino A, Esposito G, Fiorino F, et al. New Insights into the Structure-Activity Relationship and Neuroprotective Profile of Benzodiazepinone Derivatives of Neurounina-1 as Modulators of the Na(+)/Ca(2+) Exchanger Isoforms. Journal of medicinal chemistry. 2021;64(24):17901–19.

40. Wu G, Xie X, Lu ZH, and Ledeen RW. Sodium-calcium exchanger complexed with GM1 ganglioside in nuclear membrane transfers calcium from nucleoplasm to endoplasmic reticulum. Proc Natl Acad Sci U S A. 2009;106(26):10829–34.

41. Nava MM, Miroshnikova YA, Biggs LC, Whitefield DB, Metge F, Boucas J, et al. Heterochromatin-Driven Nuclear Softening Protects the Genome against Mechanical Stress-Induced Damage. Cell. 2020;181(4):800–17 e22.

42. Bickmore WA, and van Steensel B. Genome architecture: domain organization of interphase chromosomes. Cell. 2013;152(6):1270–84.

43. West G, Gullmets J, Virtanen L, Li SP, Keinanen A, Shimi T, et al. Deleterious assembly of the lamin A/C mutant p.S143P causes ER stress in familial dilated cardiomyopathy. J Cell Sci. 2016;129(14):2732-43.

44. Tasca G, D’Amico A, Monforte M, Nadaj-Pakleza A, Vialle M, Fattori F, et al. Muscle imaging in patients with tubular aggregate myopathy caused by mutations in STIM1. Neuromuscul Disord. 2015;25(11):898–903.

45. Barone V, Del Re V, Gamberucci A, Polverino V, Galli L, Rossi D, et al. Identification and characterization of three novel mutations in the CASQ1 gene in four patients with tubular aggregate myopathy. Hum Mutat. 2017;38(12):1761–73.

46. Bulla M, Gyimesi G, Kim JH, Bhardwaj R, Hediger MA, Frieden M, et al. ORAI1 channel gating and selectivity is differentially altered by natural mutations in the first or third transmembrane domain. J Physiol. 2019;597(2):561–82.

47. Endo Y, Noguchi S, Hara Y, Hayashi YK, Motomura K, Miyatake S, et al. Dominant mutations in ORAI1 cause tubular aggregate myopathy with hypocalcemia via constitutive activation of store-operated Ca(2)(+) channels. Hum Mol Genet. 2015;24(3):637–48.

48. Hedberg C, Niceta M, Fattori F, Lindvall B, Ciolfi A, D’Amico A, et al. Childhood onset tubular aggregate myopathy associated with de novo STIM1 mutations. Journal of neurology. 2014;261(5):870–6.

49. Markello T, Chen D, Kwan JY, Horkayne-Szakaly I, Morrison A, Simakova O, et al. York platelet syndrome is a CRAC channelopathy due to gain-of-function mutations in STIM1. Molecular genetics and metabolism. 2015;114(3):474–82.

50. Misceo D, Holmgren A, Louch WE, Holme PA, Mizobuchi M, Morales RJ, et al. A dominant STIM1 mutation causes Stormorken syndrome. Hum Mutat. 2014;35(5):556–64.

51. Nesin V, Wiley G, Kousi M, Ong EC, Lehmann T, Nicholl DJ, et al. Activating mutations in STIM1 and ORAI1 cause overlapping syndromes of tubular myopathy and congenital miosis. Proc Natl Acad Sci U S A. 2014;111(11):4197–202.

52. Silva-Rojas R, Treves S, Jacobs H, Kessler P, Messaddeq N, Laporte J, et al. STIM1 over-activation generates a multi-systemic phenotype affecting the skeletal muscle, spleen, eye, skin, bones and immune system in mice. Hum Mol Genet. 2019;28(10):1579–93.

53. Zhao X, Weisleder N, Thornton A, Oppong Y, Campbell R, Ma J, et al. Compromised store-operated Ca(2+) entry in aged skeletal muscle. Aging Cell. 2008.

54. Edwards JN, Blackmore DG, Gilbert DF, Murphy RM, and Launikonis BS. Store-operated calcium entry remains fully functional in aged mouse skeletal muscle despite a decline in STIM1 protein expression. Aging Cell. 2011;10(4):675–85.

