## Supplementary material for "The D84G mutation in STIM1 causes nuclear envelope dysfunction and myopathy in mice": all suppl figures

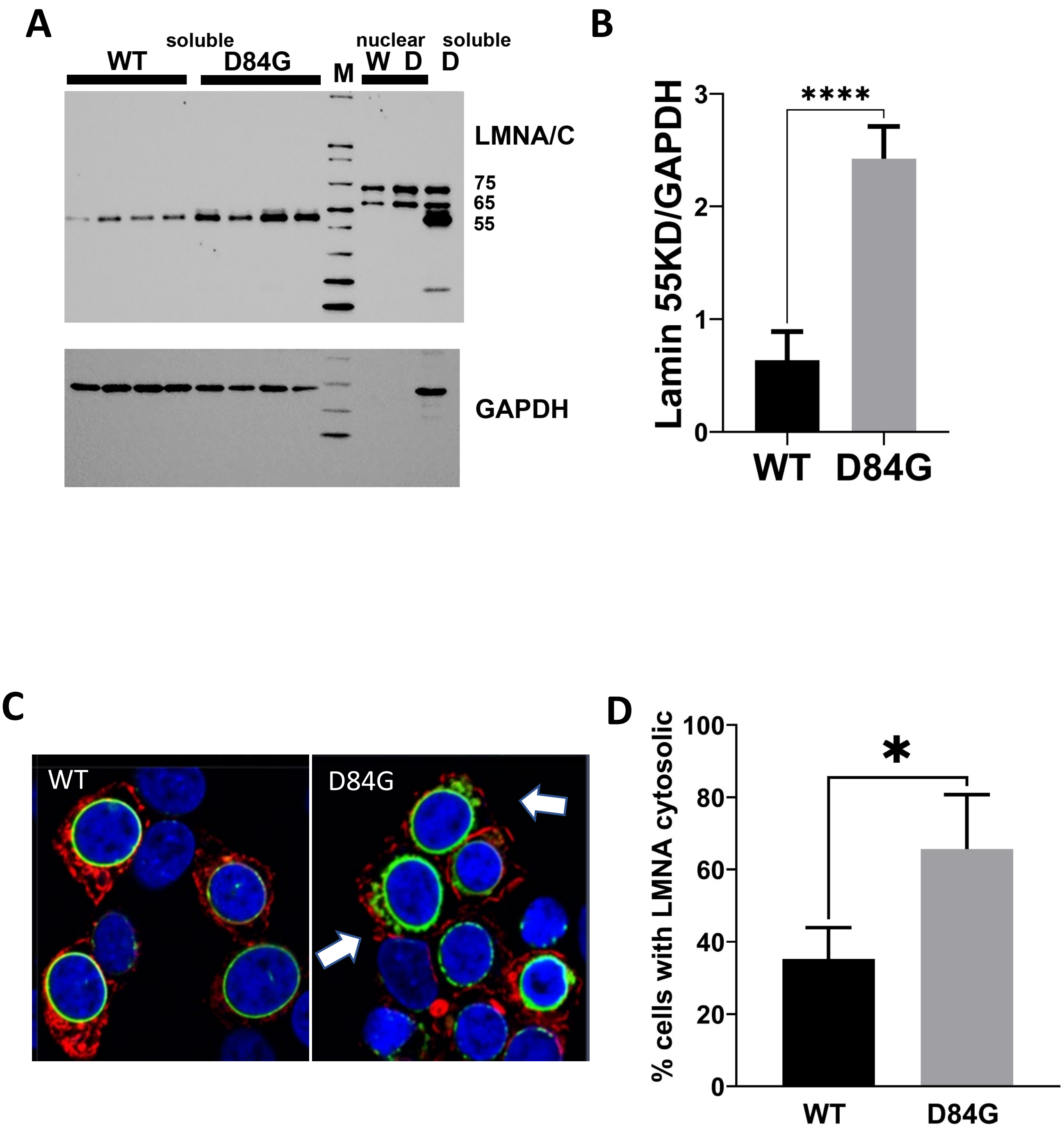


Supplemental Figure 1: D84G STIM1 damages nuclear lamina. Lysates were prepared from WT and STIM1$^{+/D84G}$ gastrocnemius muscles. Immunoblotting of the soluble and nuclear fractions detects LMNA/C as a 55kD in the soluble fraction, whereas nuclear LMNA/C is 65kD and 75kD. Lanes 1-4 are the soluble fraction from WT muscles. Lanes 5-9 are STIM1$^{+/D84G}$ muscle lysates. M identifies marker lane. Lanes 10-11 are nuclear fractions for WT (lane 10) and STIM1$^{+/D84G}$ (lane 11). Lane 12 is from lysates of STIM1$^{+/D84G}$ muscle prepared with RIPA buffer. GAPDH was used as a loading control for the soluble fraction. B) Quantification of the 55Kd band for muscle of 6-month old WT and STIM1$^{+/D84G}$ mice. **** denotes $p<0.001$. C) Confocal section for HEK293 cells expressing STIM1 (left, red) or the STIM1 mutant D84G (right, red) and LMNA-GFP (Green). Fixed cells were immunostained for STIM1 and LMNA/C. DAPI labeled nuclei (blue). D) Quantification of cytosolic LMNA-GFP. * denotes $p<0.01$. Data were obtained from 3 different experiments, 98 cells from WT and 90 cells from D84G.

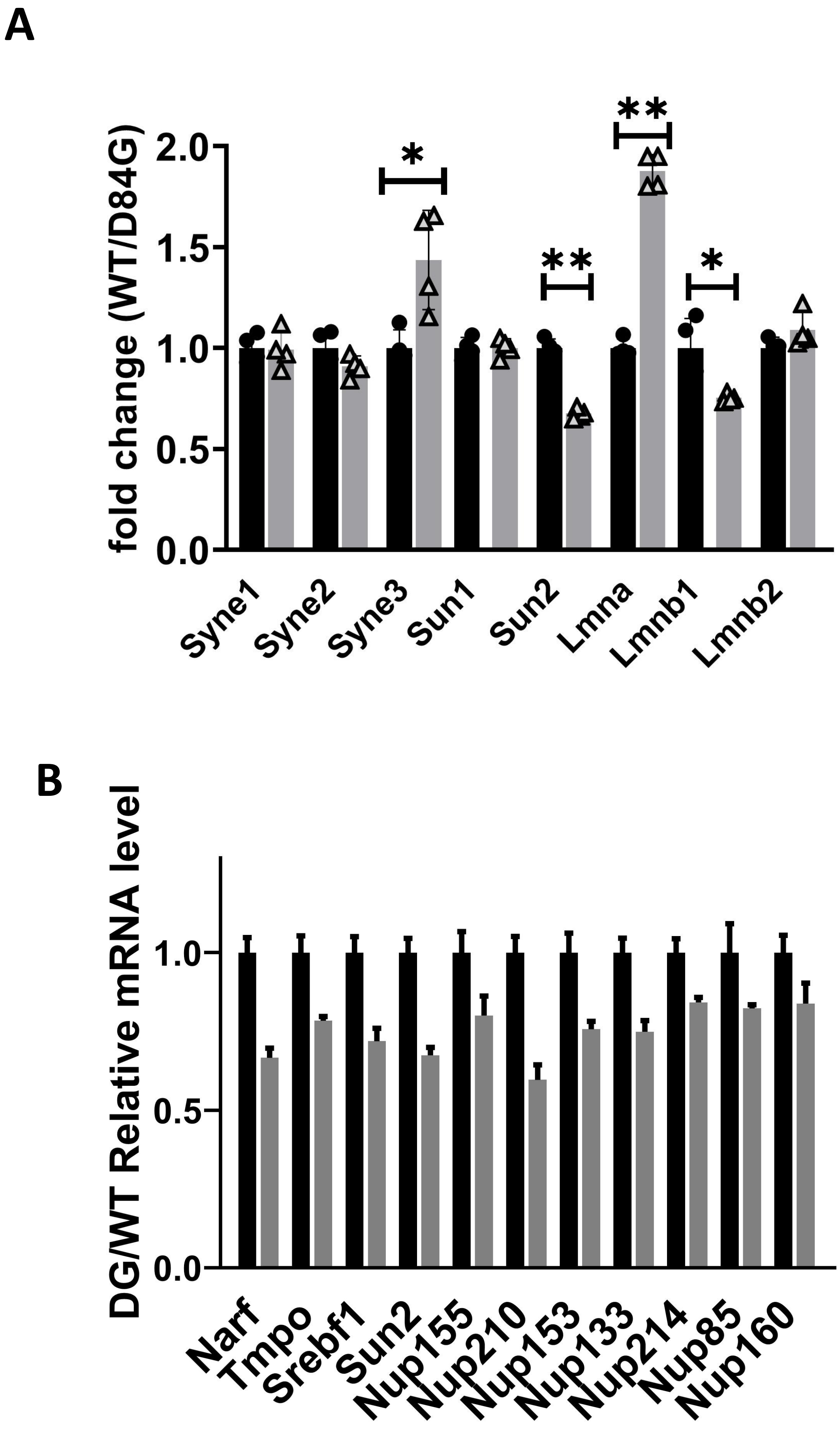


Supplemental Figure 2: Changes in mRNA expression for NE and NPC factors. A) Expression of different NE factors. Comparison of the mean ±SD for mRNA reads obtained for genes involved in nuclear lamin structure and the LINK complex. * Indicates $p<0.01$ and ** indicates $p<0.001$. n=4 mice per genotype. B) Expression of proteins associated with the nuclear pore complex (NPC). Comparison of the mean ±SD for muscles of WT and STIM1$^{+/D84G}$ mice.
